# Using cluster-based permutation tests to estimate MEG/EEG onsets: how bad is it?

**DOI:** 10.1101/2023.11.13.566864

**Authors:** Guillaume A. Rousselet

**Affiliations:** School of Psychology and Neuroscience, College of Medical, Veterinary and Life Sciences, University of Glasgow, 62 Hillhead Street, G12 8QB, Glasgow, UK

**Keywords:** Cluster inference, MEG, EEG, permutation, onset estimation, Monte-Carlo simulation, correction for multiple comparisons, family-wise error rate, false discovery rate

## Abstract

Localising effects in space, time and other dimensions is a fundamental goal of magneto- and electro-encephalography (EEG) research. A popular exploratory approach applies mass-univariate statistics followed by cluster-sum inferences, an effective way to correct for multiple comparisons while preserving high statistical power by pooling together neighbouring effects. Yet, these cluster-based methods have an important limitation: each cluster is associated with a unique *p* value, such that there is no error control at individual time points, and one must be cautious about interpreting when and where effects start and end. Sassenhagen & Draschkow (2019) provided an important reminder of this limitation. They also reported results from a simulation, suggesting that onsets estimated from EEG data are both positively biased and very variable. However, the simulation lacked comparisons to other methods. Here I report such comparisons in a new simulation, replicating the positive bias of the cluster-sum method, but also demonstrating that it performs relatively well, in terms of bias and variability, compared to other methods that provide point-wise *p* values: two methods that control the false discovery rate, and two methods that control the family-wise error rate (cluster-depth and maximum statistic methods). I also present several strategies to reduce estimation bias, including group calibration, group comparison, and using binary segmentation, a simple change point detection algorithm that outperformed mass-univariate methods in simulations. Finally, I demonstrate how to generate onset hierarchical bootstrap confidence intervals that integrate variability over trials and participants, a substantial improvement over standard group approaches that ignore measurement uncertainty.

## 1. Introduction

To localise effects in magnetoencephalography (MEG) and electroencephalography (EEG), one must apply a threshold to statistical maps. A popular frequentist approach consists in deriving a univariate threshold at every testing point, followed by a correction for multiple comparisons, often with the goal to keep at the nominal level the family-wise error rate (FWER), the probability to make at least one false positive (one type I error). A popular way to achieve this type of error control is to use cluster-based inferences, a family of methods that aim to correct for multiple comparisons while preserving high power when effects are distributed over time, space and other dimensions (Fields & Kuperberg, 2020; Frossard & Renaud, 2022; Groppe et al., 2011a, 2011b; Maris & Oostenveld, 2007; Pernet et al., 2015; Rosenblatt et al., 2018; Rousselet et al., 2014; Smith & Nichols, 2009)−for lighter introductions to cluster methods, see Ehinger (2019) and Rousselet (2018). These methods, originally introduced for structural MRI research (Bullmore et al., 1999), capitalise on the correlation of effects across measurements to control the FWER: neighbouring effects that pass a univariate threshold are lumped together, often by summing them (cluster-sum, typically of *t* or *F* values), and such combinations of effects are then compared to a reference distribution, typically a null distribution established using a permutation or a bootstrap-t approach (Pernet et al., 2015; Rousselet et al., 2023). In each permutation or bootstrap iteration, trials are randomly sampled, keeping time points (and electrodes) together, statistical tests are applied, and the resulting *t* or *F* values that exceed a threshold are grouped into clusters based on contiguity; then the maximum cluster-sum (or other cluster statistic) is saved, and the process repeated many times. Finally, a quantile (typically 95^th^ quantile) of that permutation or bootstrap distribution of maximum cluster-sums is used to assess the statistical significance of each cluster in the original data. These cluster-based techniques are particularly popular in fMRI, EEG and MEG research (for instance, as of September 2024, according to Google Scholar, Smith & Nichols (2009) has been cited over 5300 times, Maris & Oostenveld (2007) over 7800 times). Cluster-based methods also have the potential to be applied to a wider array of problems in neuroscience and other fields, as long as measurements are ordinal (Dugué et al., 2011; Rousselet, 2018).

However, cluster-based methods are not without issues. Most notably, Sassenhagen & Draschkow (2019) reminded the community of the limitations in the interpretation of cluster-based inferences and suggested guidelines on how to report such results. This useful reminder has been heard, as demonstrated by over 400 citations of the article, according to Google Scholar. Essentially, the main problem with cluster-based methods is that they provide *p* values adjusted for multiple comparisons at the cluster level only, or even for a collection of clusters for hierarchical methods such as Threshold Free Cluster Enhancement (TFCE, Ehinger, 2019; Smith & Nichols, 2009). There are no *p* values at individual testing points (aside from the unadjusted mass-univariate ones), and no error control for onsets. Consequently, cluster-based methods do not provide direct inferences about the localisation of effects. Obviously, this is deeply problematic given that localisation is one of the main goals of brain imaging. For instance, localising effects, and in particular onsets of task and stimulus modulations, has been instrumental to constrain models of visual processing, to derive a timeline of events from early sensory encoding to decision making (see for instance Bieniek et al., 2016; Foxe & Simpson, 2002; Jaworska et al., 2020; Philiastides et al., 2006; Thorpe & Fabre-Thorpe, 2001). So given the popularity of cluster-based methods, it is essential to determine if cluster-based localisation can be trusted. More specifically, how noisy are onsets estimated using cluster-based inferences? While several studies have looked at false positives and power associated with cluster-based methods (see references in the first paragraph), there is surprisingly very little research about their ability to estimate onsets.

To address this question, Sassenhagen & Draschkow (2019) reported results from the simulation of a single participant dataset, which suggests that onset estimation using cluster-based inferences is very noisy. They used a truncated Gaussian embedded in a sequence of zeros as a template, to define a real onset to be estimated. Then, they compared 50 trials of noise only, to 50 trials of noise + template, which were compared at every testing point using *F* statistics. The noise was derived from the EEG recording of one participant, and the effect was localised over a subset of time points and electrodes. Correction for multiple comparisons was achieved by using cluster-sum statistics: neighbouring *F* values that passed a threshold (*p* < 0.05) were summed, and these cluster-sums were compared to a permutation (null) distribution of maximum cluster-sums to determine statistical significance. For each simulation iteration, the first statistically significant point was declared the effect onset. The distribution of estimated onsets across 10,000 simulation iterations revealed two important patterns: estimated onsets were positively biased, with the distribution mode higher than the real onset; the distribution was asymmetric, with a thicker left tail than right tail. Sassenhagen & Draschkow (2019) reported that “on >20% of runs, the effect onset was estimated too early; divergence of 40 ms or more were found at >10% of runs.” No bias estimate was reported, even though this was a crucial aspect of the sampling distribution. These results suggest both a high risk of under-estimating onsets using cluster-based inferences, and a general tendency to over-estimate them—this superficial contradiction is explained by a skewed and heavy-tailed sampling distribution (see Figure 1 in Sassenhagen & Draschkow, 2019). However, it is difficult to fully appreciate the simulation results, because no other method was used to provide context. So here, my main goal was to estimate onsets from simulated data using other approaches, to help put the results in perspective.

**Figure 1.**
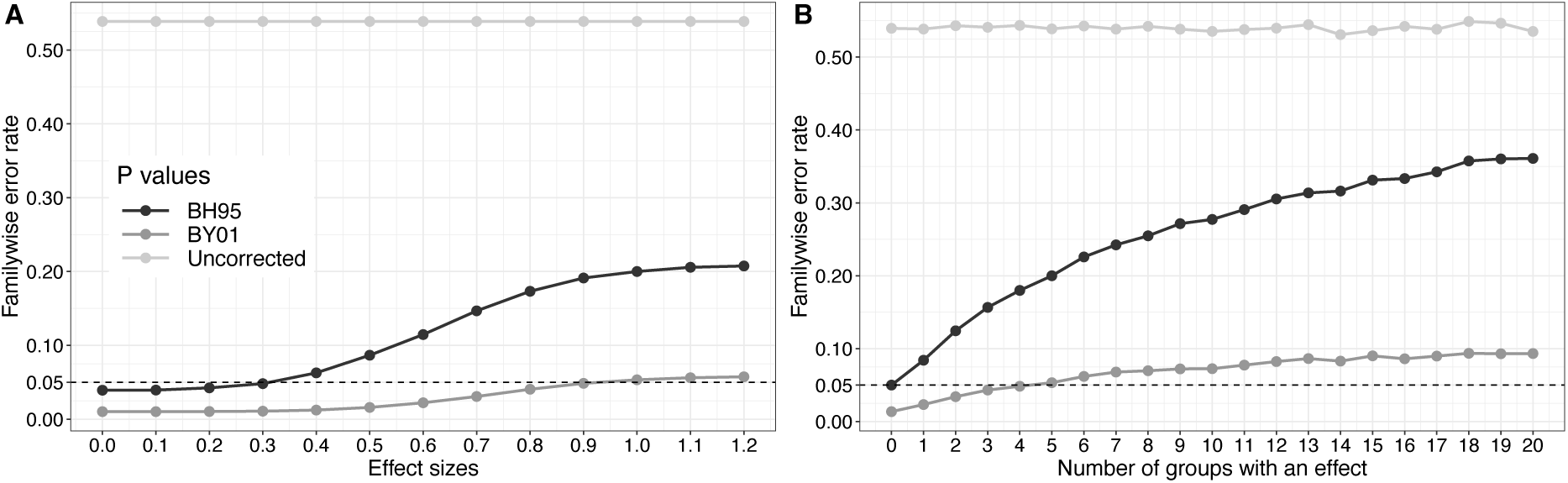
Weak control of the FWER using an FDR correction for multiple comparisons. The two panels show results of simulations with 10,000 iterations, and independent groups of 20 observations sampled from normal distributions with standard deviation of 1. A one-sample t-test was applied to the data from each group. The FWER was calculated using *p* values that were uncorrected, or corrected using either the BH95 (Benjamini & Hochberg, 1995) or the BY01 (Benjamini & Yekutieli, 2001) FDR correction. (A) Simulation with 20 groups: 15 groups had a mean of zero (no effect), and 5 groups had a mean that varies from 0 to 1.2. (B) Simulation with varying number of groups: 15 groups had a mean of zero, and 0 to 20 groups had a mean of 1. The uncorrected FWER estimates are very close to the roughly 0.54 expected in theory (1 − (1 − *alpha*)^15^). This figure can be reproduced by using the R notebook *fdr_demo.Rmd, available on GitHub* https://github.com/GRousselet/onsetsim.

When considering alternatives to cluster methods to correct for multiple comparisons, a popular approach is the false discovery rate (FDR) correction, which attempts to control the number of false positives among positive tests (Benjamini & Hochberg, 1995). Here is an example in which FDR was used specifically to address the limitations of cluster-based inferences:

“For all analyses, we corrected for multiple comparisons using a False Discovery Rate (FDR) correction (q =0.05) on standard t-tests (p <0.05), using a test that is guaranteed to be accurate for any test dependency structure, as described in Benjamini and Yekutieli (2001). This test does not suffer from some of the problems that common cluster-based permutation tests have (Maris and Oostenveld, 2007), as these may have inaccurate onset and offset boundaries (either in the temporal and/or spatial domain) due to stochastic variation in the noise (Sassenhagen and Draschkow, 2019). Note however, that prior to peer review we had initially analyzed the data using cluster-based permutation and that these analyses showed qualitatively similar results.” (van Driel et al., 2021).

The method by Benjamini & Yekutieli (2001) is a generalisation of the classic FDR correction method originally proposed by Benjamini & Hochberg (1995), which works like this, given *m p* values and a target false positive error rate α:

[1] Sort the *p* values in ascending order from 1 to *m*.
[2] Find the largest *k* for which 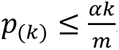.
[3] Declare statistically significant all tests associated with *p* values 1 to *k*.

For instance, if we have 10 *p* values, the first one (smallest one), *p*_(_*_k_*_=1)_ is compared to 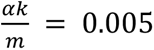, assuming α=0.05, the second one *p*_(_*_k_*_=2)_ is compared to 0.01, and so on. For the largest *p* value, *p*_(_*_k_*_=m)_ is compared to α. Once we identify *k*, we reject all tests with *p* values equal or inferior to *p*_(_*_k_*_)_. In the procedure by (Benjamini & Yekutieli, 2001), in step [2], *m* is divided by 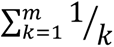, leading to a more conservative cutoff. When there is no effect, an FDR correction is equivalent to controlling the FWER; however, when many effects are expected, as is routinely the case in some research fields, including brain imaging, the FDR correction is more powerful than methods that control the FWER (Genovese et al., 2002; Nichols & Hayasaka, 2003; Winkler et al., 2024). Because of their useful properties, the 1995 and the 2001 FDR methods are available in major EEG and MEG toolboxes, such as EEGLAB (Delorme & Makeig, 2004), Fieldtrip (Oostenveld et al., 2010) and MNE python (Gramfort et al., 2013), and their application to event-related potential (ERP) data has been validated in several publications (Fields & Kuperberg, 2020; Groppe et al., 2011b, 2011a; Lage-Castellanos et al., 2010; Sheu et al., 2016). Presumably, van Driel et al. (2021), quoted above, were instructed by reviewers to use the FDR as an alternative to cluster-sum because it provides *p* values at every time point, while aiming to find the right balance between controlling false positives and maintaining high power. However, the choice of FDR was misguided because the availability of individual *p* values is not sufficient to enable localisation: in fact, it is well established that FDR methods allow limited claims about the location of effects (Genovese et al., 2002; Winkler et al., 2024). What matters is the nature of the FWER control: weak or strong (Proschan & Brittain, 2020). Weak FWER control means that the FWER is at most at the alpha level only when the global (omnibus) null hypothesis is true (all null hypotheses are true). When we reject under weak control, we reject the omnibus hypothesis and conclude that the results suggest the presence of an effect somewhere, but strict localisation is not possible. Cluster-based and FDR methods offer weak FWER control. In the case of FDR, weak control is due to the presence of effects changing the overall distribution of *p* values, which can lead to an excess of *p* values associated with true null hypotheses to be considered significant. This in turn leads to a FWER higher than the nominal level. So standard FDR methods control the error rate globally, not for any subset of tests, for instance in separate time-windows, or separately for clusters of positive or negative group differences. For instance, when applying two-sided tests, false discovery rates can be dramatically inflated for test statistics in the direction opposite to that of the signal (Winkler et al. 2024).

The weak FWER control afforded by FDR is worth illustrating, given that this limitation is often ignored by researchers. In Figure 1, I show results from simple simulations (like all figures and results in this article, Figure 1 can be reproduced by using the code mentioned in the caption). In panel A, we look at the FWER across 15 groups for which there is no effect, as a function of the results in 5 groups in which the effect sizes vary from 0 to 1.2. As is evident, even though the FDR methods provide some control over the FWER (it is much lower than without correction), it can be much higher than the nominal alpha level (here 0.05), and it grows with effect sizes in other parts of the data. Similarly, in panel B, the FWER grows with the number of groups with a fixed effect size. In both panels, it is also apparent that the BY01 method (Benjamini & Yekutieli, 2001) is much more conservative than the original BH95 method (Benjamini & Hochberg, 1995; see details with clear illustrations in Winkler et al., 2024).

In contrast to cluster-based and FDR methods, which offer weak control of the FWER, Bonferroni and related methods offer strong FWER control (Hochberg, 1988). However, these methods are too conservative to be of any use when testing the very large number of hypotheses typically encountered in brain imaging research. A useful alternative is the maximum statistics correction for multiple comparisons (Groppe et al., 2011a, 2011b; Holmes et al., 1996; Ince et al., 2017; Nichols & Holmes, 2002). The MAX correction achieves strong control of the FWER by comparing each statistic, say a *t* or *F* value, to a distribution of maximum statistics obtained under the null: for each permutation or bootstrap iteration, save the maximum absolute statistics across all tests; repeat many times, say 2,000 times; then use a quantile of the distribution of maximum statistics as a threshold for all tests performed on the original data. MAX corrected statistical maps can be used to localise effects, at the cost of lower power, and are thus worth considering as an onset detection method.

Another method worth considering (and suggested by one of the reviewers), is the more recent and very promising cluster-depth approach, which uses cluster inferences, while delivering strong control of the FWER (Frossard & Renaud, 2021, 2022). The method works by forming clusters, but instead of making inferences on cluster statistics (like the sum of *F* values inside a cluster), it uses a permutation test to form null distributions based on the position inside the cluster: when a cluster is identified in the original data, the statistic observed at position 1 inside the cluster is evaluated against a distribution of statistics observed at position 1 in clusters obtained in permuted data, and so on for the different positions. That way, a *p* value is obtained at each time point. By default, two *p* values are calculated for each time-point within a cluster, one by using positions relative to the start of the cluster (“from the head”), another one by using positions relative to the end of the cluster (“from the tail”), and the maximum of these two *p* values is returned (see full details, proofs, and extensive validations in Frossard & Renaud, 2022).

Frossard & Renaud (2022, section 4.2) explain that to achieve strong FWER control in the cluster-depth procedure, it is required to remove the average from each group of observations, so as to obtain stationary residuals. However, it is important to clarify that this subtraction procedure is not necessary to obtain strong control (the MAX method achieves strong control without it), and it is not sufficient (FDR corrected and cluster-sum *p* values derived from permutation distributions after mean subtraction still provides weak FWER control). This point is illustrated in Figure 2 (panel A), which reports a simulation in which two independent groups were compared at each of 20 time points: data at the first 15 time points were sampled from the same population (no effect), whereas the groups at the last 5 time points differed by an effect size ranging from 0 to 1. Replicating the results from Figure 1, with FDR methods, the FWER increases with the effect size in the cluster of true effects; the same occurs with the cluster-sum technique—these methods provide weak FWER control. In contract, the FWER is relatively stable across effect sizes when using the cluster-depth and MAX methods—they offer strong control of the FWER.

**Figure 2.**
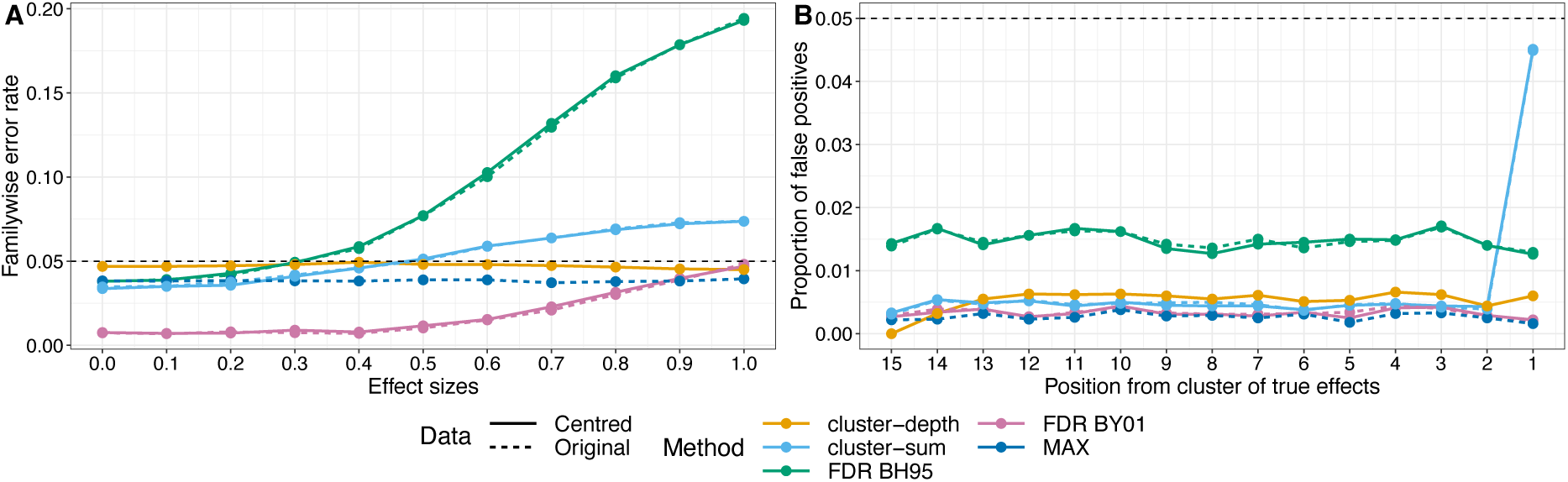
Weak and strong FWER control. Results of a simulation with 10,000 iterations. Two-sample t-tests were applied at 20 time points. At each time point, two independent groups were compared, each with 30 trials. At the first 15 time points, both groups were sampled from a population with mean 0. At the last 5 time points, one group came from a population with mean 0, the other group from a population with a mean that varied between 0 and 1. All populations were normally distributed with standard deviation 1. *P* values were obtained using a permutation test with 2,000 iterations. Permutation distributions were derived from either the original data or centred data (mean subtracted from each group). For the cluster-sum and cluster-depth, the cluster forming threshold was *p* ≤ 0.05. (A) FWER as a function of effect sizes. (B) Proportion of false positives at each of the 15 time points without effect, as a function of their position relative to the cluster of 5 time points, when groups differed by an effect size of 1 (the maximum difference considered in panel A). Position 1 is the closest to the cluster with large group differences. This figure can be reproduced by using the R notebook *weakstrongfwer_demo.Rmd*.

If we look at the proportion of false positives at each of the 15 time points at which the populations do not differ, it remains constant across locations, except for the location adjacent to the cluster of true effects (position 1 in Figure 2, panel B), and only when using cluster-sum. This result fits with the intuition that noise can be lumped together with signal inside a cluster, thus potentially distorting onset estimates. Here, the effect is limited to one time point because white noise was used. Auto-correlated noise, typical of physiological recordings, could potentially lead to positions further away from the cluster to be affected. In the case of FDR methods, the increase in false positives is not position specific, because it only depends on the ranking of *p* values.

Yet other onset estimation methods have been proposed, for instance defining the onset as the time at which a certain proportion of the peak amplitude or the cumulative peak amplitude has been reached (Fahrenfort, 2020; Liesefeld, 2018). However, such methods require that an effect has already been detected, and more importantly, they depend on potentially subjective peak identification, which, from experience, can be problematic when making inferences in individual participants. These methods are also guaranteed, by definition, to lead to positively biased onset estimates because they do not seek to identify the earliest detectable difference. Hence, these methods are not considered here but maybe they have some merit. The TFCE method is also not considered because it is necessarily more biased than the standard cluster-sum (see Figure 1 in Smith & Nichols, 2009).

Finally, all the methods considered so far have a key limitation. Whether they provide weak or strong FWER control, cluster-sum, FDR, MAX, and cluster-depth do not solve a fundamental issue: whether a *p* value is available at each testing point or not is irrelevant, because these methods do not explicitly assess onsets. That’s because onsets are second-order properties of the data. As such, an onset is defined by a sustained lack of effect, followed by the sustained presence of an effect—it is thus a multivariate interaction. A researcher seeking to establish the start of an effect using a mass-univariate approach, even when combined with a method that provides strong FWER control, commits 3 errors: (1) accepting the null based on a *p* value larger than alpha (Riesthuis, 2024), (2) interpreting a series of univariate tests as a multivariate pattern, without modelling it explicitly (Simpson, 2018), (3) declaring significant the difference between significant and non-significant effects, without testing it—an interaction fallacy (Gelman & Stern, 2006; Nieuwenhuis et al., 2011). Instead, the interaction that defines an onset is an inference performed essentially by eyeballing the results. This problem led me to consider methods that could explicitly estimate onsets in time-series, without being computationally prohibitive. For instance, there is a vast family of algorithms aimed at detecting change points in time-series (Aminikhanghahi & Cook, 2017; Chen & Gupta, 2013; Lindeløv, 2020, 2023; Truong et al., 2020; Zhao et al., 2013). Here, for comparison with the cluster-sum, FDR, MAX and cluster-depth methods, I explored the feasibility of using a simple change point detection method to estimate EEG onsets. As a proof of concept, I considered the well-established binary segmentation algorithm, which is fast to run (Aminikhanghahi & Cook, 2017; Eckley et al., 2011; Edwards & Cavalli-Sforza, 1965; Scott & Knott, 1974; Truong et al., 2020). The algorithm was applied to changes in mean and variance (Chen & Gupta, 2012), which seems appropriate to detect changes in relatively slow ERP time-courses (see an excellent illustration in figure 2 of Kappenman et al., 2021). Unlike cluster-based, FDR and MAX methods, the change point method was applied directly to the time-course of the *t* values from the independent t-tests, so it relies on a different type of inference.

In this article, I report simulation results contrasting the onset estimation performance of the above methods. This was done in the simplified situation in which we look for an onset at a single electrode, in one participant. Then I consider how individual onsets can be used to make group inferences, and the benefits of doing so. I conclude with recommendations on the implementation of an onset estimation pipeline in EEG and MEG research.

## 2. Methods

All the figures and simulations presented here can be reproduced using R code (R Core Team, 2021) available on GitHub: https://github.com/GRousselet/onsetsim. A README file contains a table listing all the main simulations and figures, with the matching R notebooks. Similar results were obtained using a Matlab version (The MathWorks Inc., 2021) of some of the simulations. Only the R results are presented here. During the review process, one of the reviewers also replicated the main simulation results (Ehinger, 2024), using an implementation in the Julia programming language (Bezanson et al., 2017). I also performed simulations using less complex 1/f type of noise (Stevens, 2009). Using pink noise gave very similar results to the ones reported here, but using white noise substantially altered the behaviour of the FDR method—see full simulations on GitHub.

### 2.1. Simulations

Following the approach used by Sassenhagen & Draschkow (2019), I compared two conditions, one with noise only, one with noise + signal. The noise was generated by superimposing 50 sinusoids at different frequencies, following an EEG like spectrum (see details and code in Yeung et al., 2004). Examples of 4 noise-only trials are presented in Figure 3A. On each trial, the noise was normalised to have a variance of 1. Increasing the variance to 1.5 affected the Monte-Carlo summary statistics of all methods, but did not affect the conclusion of the simulations. The signal was a truncated Gaussian defining an objective onset at 160 ms, a maximum at 250 ms, and an offset at 342 ms (Figure 3B). It was created by getting the probability density function of a standard normal with 93 points from x = -1.5 to 1.5, rescaled in amplitude between 0 and 1, and padded with 79 zeros to the left and to the right, to form a time-series of 251 points, from 0 to 500 ms, in steps of 2 ms (500 Hz sampling rate).

**Figure 3.**
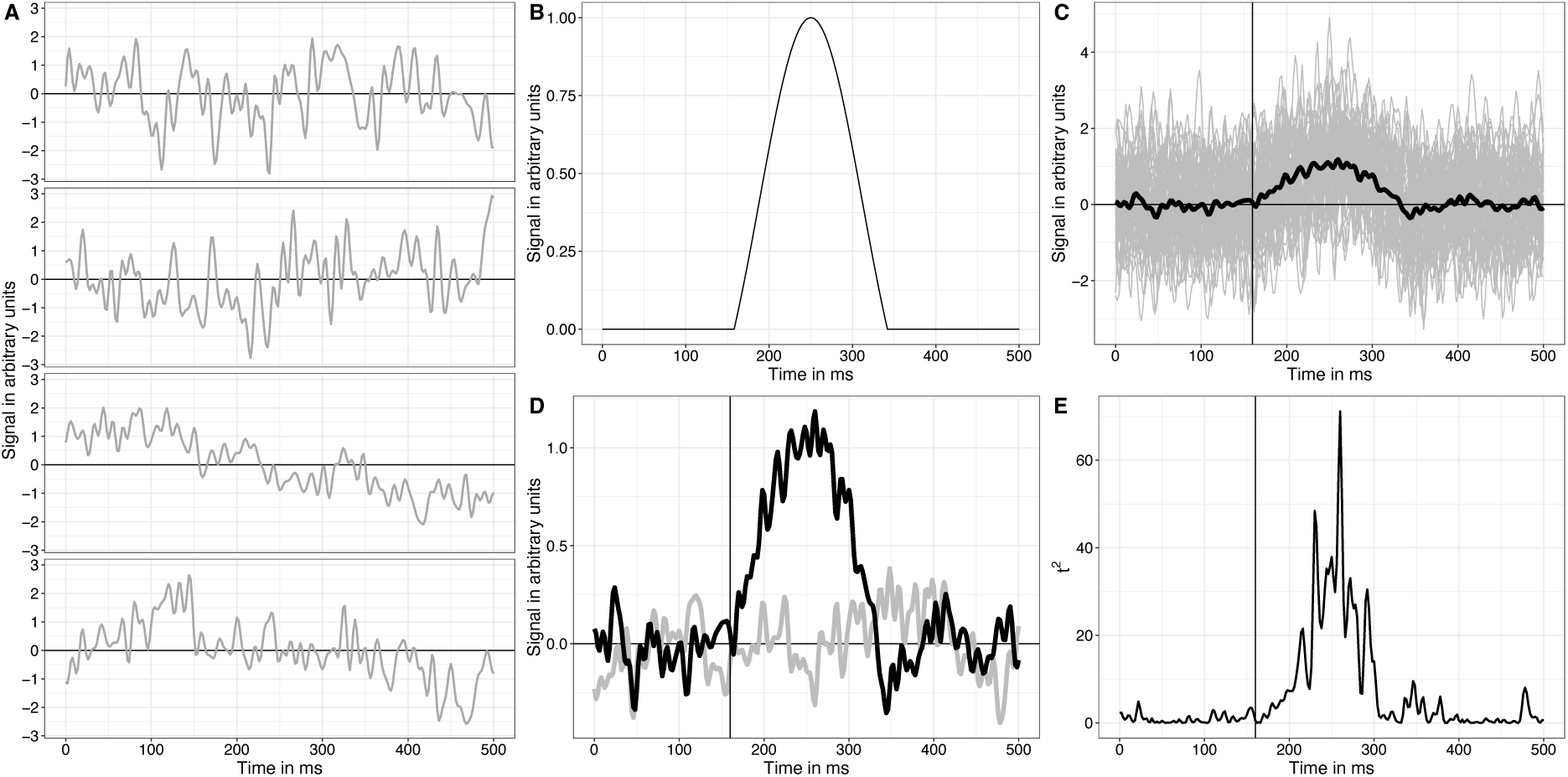
Data generation process. (A) Examples of 4 noise trials. (B) Truncated Gaussian used as signal. (C) Example of 50 trials of noise + signal (grey), with the superimposed mean (black). (D) Example of averages of 50 trials for the noise only condition (grey) and the noise + signal condition (black). (E) Time-course of squared *t* values from a series of t-tests for two independent samples with unequal variances. This figure can be reproduced by using the R notebook *examples.Rmd*.

Each Monte-Carlo simulation consisted of 10,000 iterations, starting with the creation of 50 trials of the noise only condition, and 50 trials of the noise + signal condition (Figure 3C, D). The two conditions were then compared at every time point using a t-test for two independent samples with unequal variances, and the *t* values squared (*t*^2^, Figure 3E). This approach is very similar to that of Sassenhagen & Draschkow (2019), except that I simulated only one electrode, with cluster inferences only in the time domain. There is no need to simulate extra dimensions (electrodes, time-frequencies), because the argument against cluster-based inferences is not specific to one dimension. Also, I used synthetic noise instead of using data from one participant.

To assess the accuracy of group onset estimation, I ran a simulation like the one already described, but now with 20 participants. In each simulation iteration, each participant was assigned a random onset between 150 to 170 ms, drawn from a uniform distribution with 2-ms steps. These random onsets were created by adjusting the position of the truncated Gaussian used to generate individual trials. Once the onsets were estimated for every participant, group onsets were quantified as the median onset across participants, separately for each method. Using other quantiles of the distribution of onsets is also considered in the result section.

This group example was extended to a situation in which onsets from two groups are compared. Each group was composed of 20 participants, with, for each participant, 50 noise trials and 50 signal trials generated as described above. In the first group, each participant had a random onset sampled from a uniform distribution ranging from 140 ms to 180 ms. In the second group, onsets were sampled from one of three uniform distributions: 140-180 ms (no group difference); 130-190 ms (group difference in spread); 150-190 ms (group shift).

Finally, I considered an example involving a hierarchical bootstrap approach, in which participants are sampled with replacements, followed by sampling with replacements of trials, independently in each condition (Rousselet et al., 2023). The example involved different situations, with one group of 30, 50 or 100 participants, and 30, 50 or 100 trials per condition. Individual onsets were sampled from a discrete uniform distribution 130-190 ms, in steps of 2 ms, and the amplitude of the truncated Gaussian was sampled from a uniform distribution 0.5-1, in arbitrary units. Signal and noise were otherwise generated as previously described.

### 2.2. Inferential statistics and onset estimation

Null distributions of *t^2^*values were estimated using a permutation with k=2,000 iterations (Maris & Oostenveld, 2007; Pernet et al., 2015). In each permutation iteration, trials from the two conditions were combined before being randomly assigned to each surrogate condition, keeping all time points together, and a series of *t* values calculated. At each time point, the distribution of k permutation *t*^2*^ values was used to compute univariate *p* values using the formula 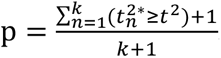. Alpha was set to an arbitrary 0.05. Five methods for multiple comparison corrections were considered: two FDR methods; two cluster-based methods; and the maximum statistics.

The two FDR methods were BH95 (Benjamini & Hochberg, 1995) and BY01 (Benjamini & Yekutieli, 2001), which were applied to the permutation *p* values using the *p.adjust* function in R.

The first cluster-based inference was implemented using a cluster-sum statistic of squared *t*^2^ values, using my own R version of the implementation in the *LIMO EEG* toolbox (Pernet, Chauveau, et al., 2011; Pernet et al., 2015). The cluster forming threshold was set to an arbitrary 0.05. In the original *t*^2^ time-series, contiguous *t*^2^ values that passed the univariate cluster-forming threshold were summed. Each of these sums was compared to the 95^th^ quantile of the distribution of maximum *t*^2*^ sums obtained in the permutation: for each iteration of the permutation, clusters were identified, *t*^2^ values summed in each cluster, and the largest sum saved, leading to a distribution of k maximum *t*^2*^ sums. I will refer to this method as cluster-sum or method CS in the results. The second cluster-based inference, cluster-depth (CD) was like the first one, except that the permutation used data that had been centred independently at each time point and for each condition (the mean was removed from individual observation), and a *p* value was not calculated for the entire cluster, but for each time-point inside the cluster, based on its position relative to the start of the cluster (Frossard & Renaud, 2022). This was done using the *clusterlm* function, from the *permuco* R package (Frossard & Renaud, 2021), with the argument *multcomp* set to "clusterdepth_head". This option is less conservative than the default, which consists in returning the maximum between the *p* value calculated from the head and the one calculated from the tail. As only onsets are of interest here, it seems appropriate to accept a slightly increased type I error rate (false positives) to improve our sensitivity to early effects.

The maximum statistics (MAX) correction (Holmes et al., 1996; Nichols & Holmes, 2002) was applied by using a unique threshold for all *t*^2^ values, the 95^th^ quantile of the permutation distribution of maximum *t^2^* values: in each permutation iteration, the maximum *t*^2*^ value was saved, leading to a distribution of k maximum *t*^2*^ values. For the cluster-sum, FDR and MAX methods, the onset was defined as the first statistically significant time point after correction for multiple comparisons. However, I discarded the first cluster of significant points if it included zero, as this is unrealistic (Bieniek et al., 2016). Including onsets at zero ms was only detrimental to the FDR method.

Change point detection was applied to the time-course of *t^2^* values using the binary segmentation algorithm (Edwards & Cavalli-Sforza, 1965; Scott & Knott, 1974), which was set to look for changes in mean and variance of normally distributed data (Chen & Gupta, 2012). The penalty applied to the likelihood cost function was the modified Bayes information criterion (MBIC, Zhang & Siegmund, 2007) and the minimum segment length was two. The number of expected change points was set to two and the first of these two change points was recorded as the effect onset. This approach is implemented in the *cpt.meanvar* function from the *changepoint* R package (Killick et al., 2022; Killick & Eckley, 2014). Other parameters were left to their default values. Using more than two expected change points would lead to higher variability in onset estimation, thus this approach provides a good benchmark about what can be achieved with a simple change detection algorithm, assuming we already know that an effect exists.

Figure 4 shows the onsets estimated by each of the six methods, using the data from the example in Figure 3. In grey is the permutation distribution of *t*^2^ values. The solid vertical line indicates the true onset (160 ms). The dotted vertical line marks the estimated onset. The horizontal dashed lines are univariate permutation thresholds.

**Figure 4.**
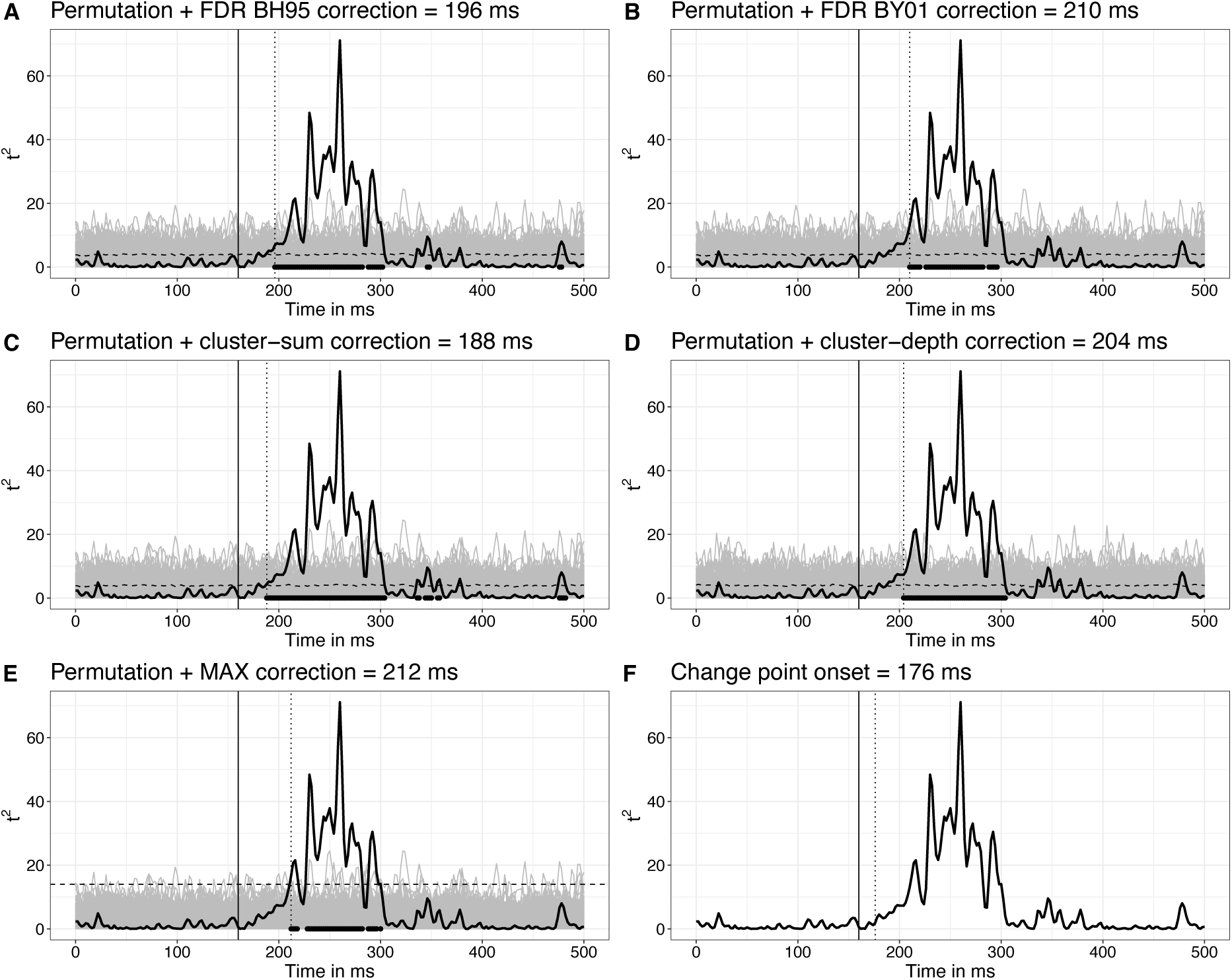
Examples of onset inferences. The black vertical lines indicate the true onset (160 ms). The dotted vertical lines mark the estimated onsets. Each panel shows the same time-course of *t^2^* values in black. The superimposed grey lines are the 2,000 *t^2*^* values from the permutation. Dashed horizontal lines mark the 1-alpha quantiles of the univariate permutation distributions. (A-B) The permutation distributions were used to compute a *p* value at each time point, before applying the FDR correction. (C) The dashed line is the same as in panel A. Neighbouring points above these univariate thresholds were clustered, and for each cluster the sum of the *t^2^* values was assessed against the 1-alpha quantile of the permutation distribution of maximum cluster-sums. (D) The cluster-depth algorithm also relied on a permutation distribution, but unlike cluster-sum, it returns a *p* value at each location in the cluster. (E) For each permutation, the maximum value was saved, and the 1-alpha quantile of the distribution of maxima is illustrated as a straight dashed line. (F) The change point algorithm was applied directly to the time-course of *t^2^* values, without considering the permutation distribution. This figure can be reproduced by using the R notebook *examples.Rmd*.

### 2.3. Quantification of onset results

Onset estimation methods were assessed using these summary statistics of the Monte-Carlo sampling distributions of onsets: bias, mean absolute error (MAE) and variance (Aminikhanghahi & Cook, 2017; Morris et al., 2019). Bias was quantified by subtracting the true onset from the median of each sampling distribution. Because the distributions are asymmetric, the median provides a better reflection of the typical onset than the mean (Rousselet & Wilcox, 2020). The MAE was the average of the absolute differences between onset estimates and the true onset. Following Sassenhagen & Draschkow (2019), I also report the proportion of estimated onsets that occurred before the true onset (proportion too early).

## 3. Results

After 10,000 iterations, for our six methods, we get the distributions of estimated onsets in Figure 5A. Each distribution has a shape similar to that reported by Sassenhagen & Draschkow (2019): overall a positive bias, with the modes shifted to right of the true onset (black vertical line), and some amount of under-estimation, which is particularly visible for the FDR BH95 and change point methods.

**Figure 5.**
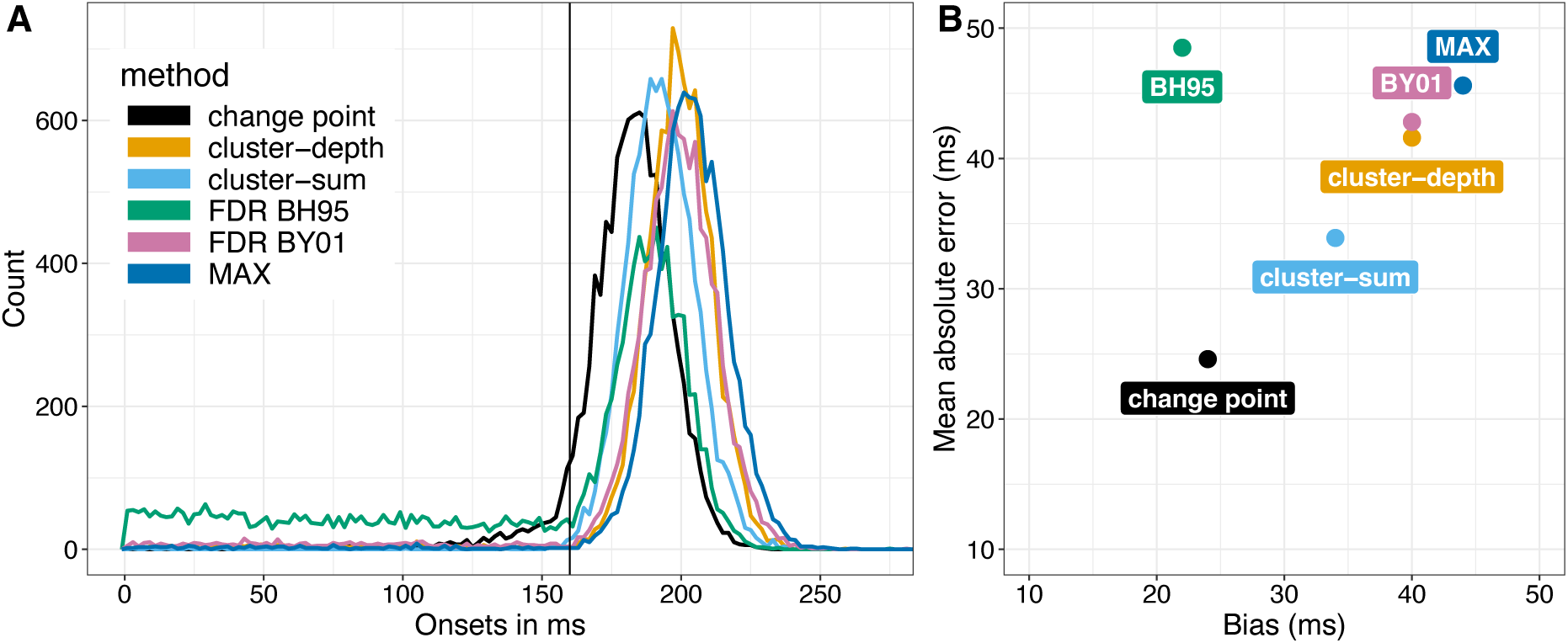
Onset sampling distributions and performance statistics. (A) Sampling distributions of estimated onsets. The vertical black line indicates the true onset (160 ms). (B) Summary statistics. This figure can be reproduced by using the R notebook *onsetsim_eeg.Rmd*. A version of panel A with smoother distributions based on 200,000 onsets for each method is available in the notebook *onsetsim_eeg_group.Rmd*.

We can quantify the results by considering summary statistics of these distributions: bias, mean absolute error (MAE), variance, and proportion too early (before the true onset). These summary statistics are reported in Table 1, and the mean absolute error is plotted as a function of bias in Figure 5B.

**Table 1.** Summary statistics of the sampling distributions presented in Figure 5. Methods: CP = change point; CD = cluster-depth; CS = cluster-sum; FDR BH95 = false discovery rate using the Benjamini-Hochberg (1995) method; FDR BY01 = false discovery rate using the Benjamini-Yekutieli (2001) method; MAX = maximum statistics; MAE = mean absolute error. The content of this table can be reproduced by using the R notebook *onsetsim_eeg.Rmd*.

| Statistic | CP | CD | CS | FDR BH95 | FDR BY01 | MAX |
| --- | --- | --- | --- | --- | --- | --- |
| **Bias (ms)** | 24 | 40 | 34 | 22 | 40 | 44 |
| **MAE (ms)** | 24.8 | 41.5 | 34 | 48.6 | 42.5 | 45.8 |
| **Variance (ms<sup>2</sup>)** | 263 | 582 | 166 | 3777 | 981 | 447 |
| **% too early** | 5.6 | 3 | 0.3 | 32.3 | 5.4 | 1.5 |

Confirming our visual inspection of the distributions, all methods are positively biased: the least biased were FDR BH95 and change point, the most biased was MAX; cluster-sum fell in between; cluster-depth and FDR BY01 were almost as biased as MAX, and more so than cluster-sum. The high bias of MAX is not surprising, as it is conservative, the price to pay for strong FWER control. The slightly higher bias of cluster-depth relative to cluster-sum is a new result, but was anticipated given the example reported in Figure 6 of Frossard & Renaud (2022). The high bias of FDR BY01 is in keeping with the literature, showing that it is much more conservative than FDR BH95 (Groppe et al., 2011b). The lower bias of FDR BH95 is however deceptive: the mode of the BH95 onset distribution (192 ms) is close to that of cluster-sum (190 ms), but the FDR BH95 distribution also has a very thick left tail. As a result, FDR BH95 is associated with the largest mean absolute error (MAE) and the largest variance. In contrast, cluster-sum had the second lowest MAE after change point, and the lowest variance among the six methods. In comparison, FDR BH95 had a variance 23 times larger than cluster-sum. As Sassenhagen & Draschkow (2019) did, if we consider the proportion of onsets earlier than the true onset, cluster-sum outperformed the other methods.

**Figure 6.**
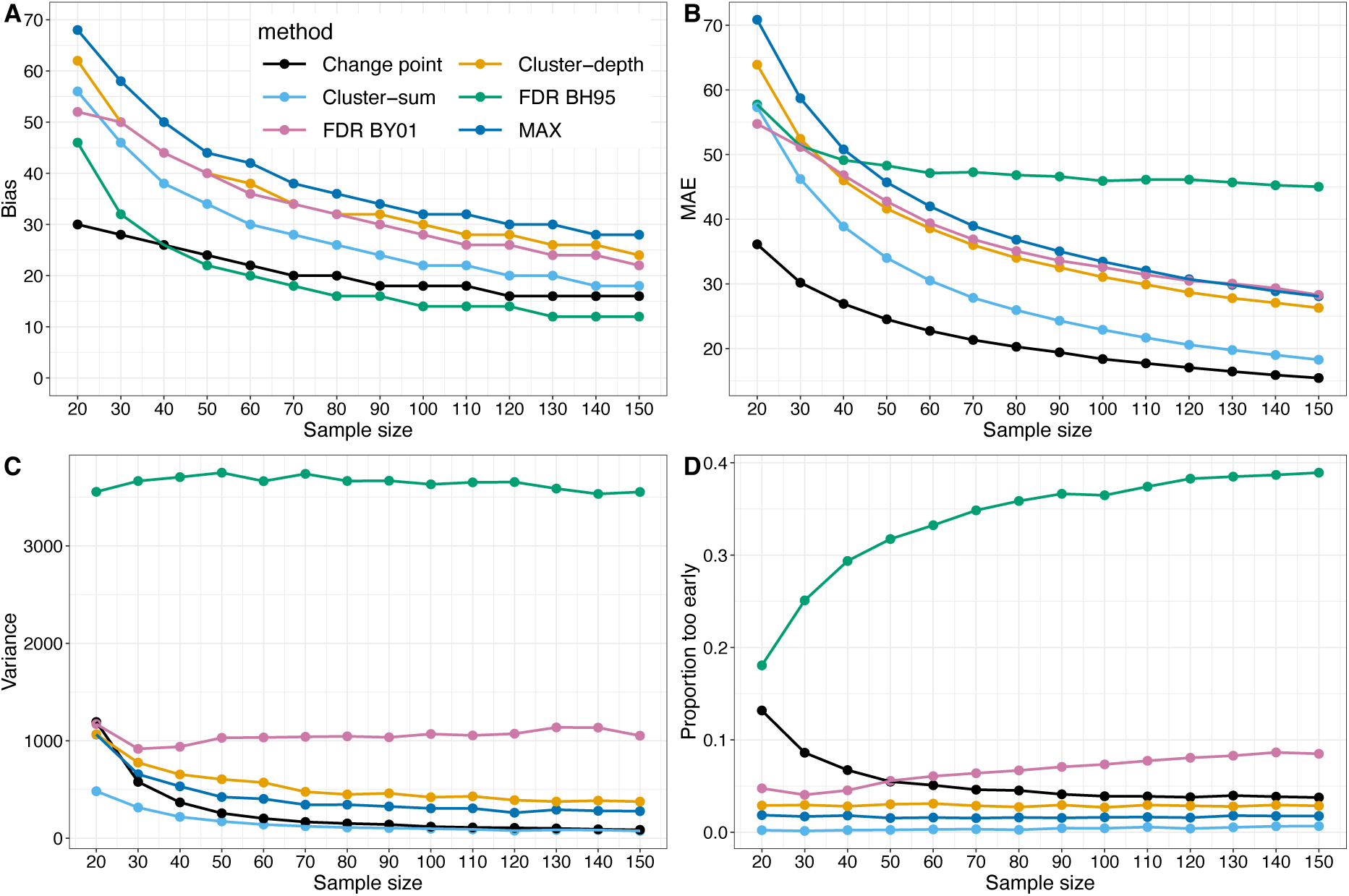
Summary statistics as a function of sample size. (A) Bias. (B) Mean absolute error (MAE). (C) Variance. (D) Proportion of estimated onsets that occurred before the true onset. This figure can be reproduced by using the R notebook *onsetsim_eeg.Rmd*.

These results are not specific to a sample size of 50 trials per condition, as demonstrated in Figure 6, which presents results for sample sizes ranging from 20 to 150 trials. Bias can be reduced considerably by using larger sample sizes, but the MAX method remains overly conservative (Figure 6A). MAE, variance and proportion of early onsets all improved with increasing sample size too, except for FDR methods (panels C and D). Again, cluster-sum performed relatively well: despite somewhat high bias (still 18 ms for n=150), it had lower MAE than the two FDR and the MAX methods, and the lowest variance and proportion of too early onsets of all methods. The change point approach looks very promising, with relatively low bias, MAE and variance, but higher under-estimation than MAX and cluster-based inferences.

As another partial check on generalisability, I used a truncated Gaussian half the width of the one used in the main simulations, keeping the noise constant: the ranking of methods was very similar, with the change point results having the lowest bias and MAE of all methods, and second lowest variance after cluster-sum (see notebook *onsetsim_eeg_half.Rmd*).

So far, we have considered results from one simulated participant. What happens in a more realistic situation in which we try to estimate onsets from 20 participants instead of one? To find out, let’s assume that each participant has a random onset between 150 and 170 ms, and 50 trials per condition. In each simulation iteration, after estimating the onset in each of the 20 participants, I used the median onset as the group estimate. This leads to the sampling distributions of estimated group onsets in Figure 7.

**Figure 7.**
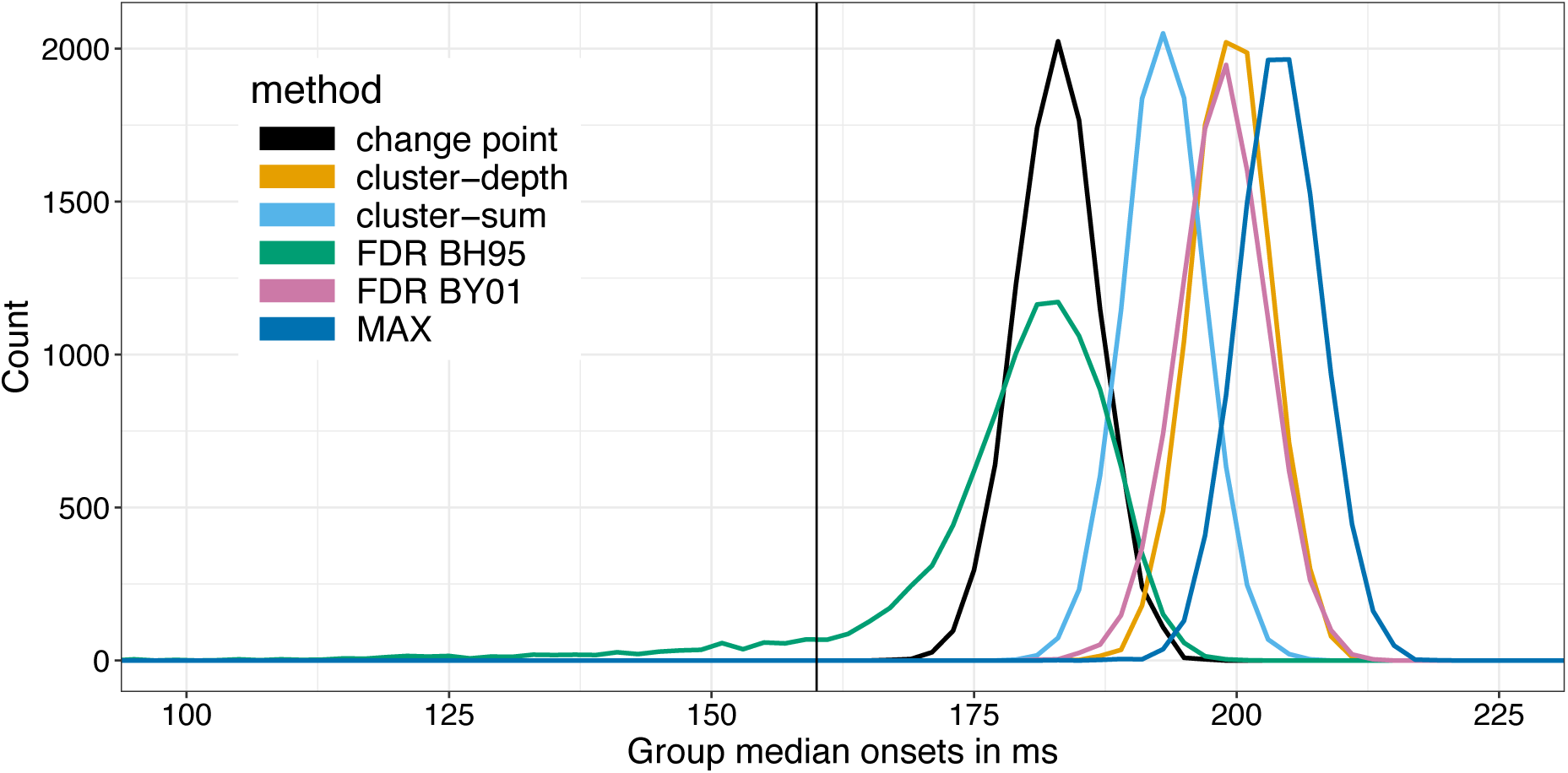
Sampling distributions of estimated group onsets. The vertical black line indicates the true onset (160 ms). This figure can be reproduced by using the R notebook *onsetsim_eeg_group.Rmd*.

The group results clearly distinguish the six methods, with, in increasing order of bias and MAE: BH95, change point, cluster-sum, BY01, cluster-depth and MAX (Table 2). Variance is again dramatically larger for the BH95 method than for the other methods, which do not differ much from each other. BH95 is also the only method that produces group under-estimation. For the other methods, combining onsets using the median across participants eliminates under-estimation.

**Table 2.** Summary statistics of the sampling distributions presented in Figure 7. Methods: CP = change point; CD = cluster-depth; CS = cluster-sum; FDR BH95 = false discovery rate using the Benjamini-Hochberg (1995) method; FDR BY01 = false discovery rate using the Benjamini-Yekutieli (2001) method; MAX = maximum statistics. MAE = mean absolute error. The content of this table can be reproduced by using the R notebook *onsetsim_eeg_group.Rmd*.

| Statistic | CP | CD | CS | FDR BH95 | FDR BY01 | MAX |
| --- | --- | --- | --- | --- | --- | --- |
| **Bias (ms)** | 24 | 40 | 34 | 21 | 39 | 45 |
| **MAE (ms)** | 23.6 | 39.9 | 33.6 | 20.9 | 39.4 | 44.7 |
| **Variance (ms<sup>2</sup>)** | 16 | 14 | 14 | 145 | 18 | 15 |
| **% too early** | 0 | 0 | 0 | 5.7 | 0 | 0 |

Unsurprisingly, the distribution of group median onsets is less noisy than the distribution of individual estimates, but it remains positively biased. We can try to bring these group estimates closer to the population onset value by exploring bias as a function of the quantiles of onsets, as illustrated in Figure 8A. The idea is to calibrate the group quantile to obtain the minimum positive group bias for each method. In this simulation, based on the results in Figure 8A, the solution is to use the 0.1 quantile for the BY01 and change point methods; the 0.35 quantile for BH95; and the 0.05 quantile for the other methods. The matching sampling distribution of group onsets are illustrated in Figure 8B. This approach works remarkably well for the change point method, with a sampling distribution almost centred on the true onset. For the other methods, while bias can be reduced, the new distributions have very long left tails, which means that group onsets would underestimate the true onsets in many experiments. This is particularly problematic for the most biased methods, cluster-sum, cluster-depth, and MAX, because the more extreme the quantile, the noisier its estimation (Wilcox & Rousselet, 2024). Adding more trials and participants would improve estimation of extreme quantiles, but it might be prudent to consider higher quantiles for improved accuracy, even if that means higher bias. Obviously, such an approach should be validated for specific applications, ideally using simulations to determine the sample sizes required to achieve certain levels of statistical power and measurement precision.

**Figure 8.**
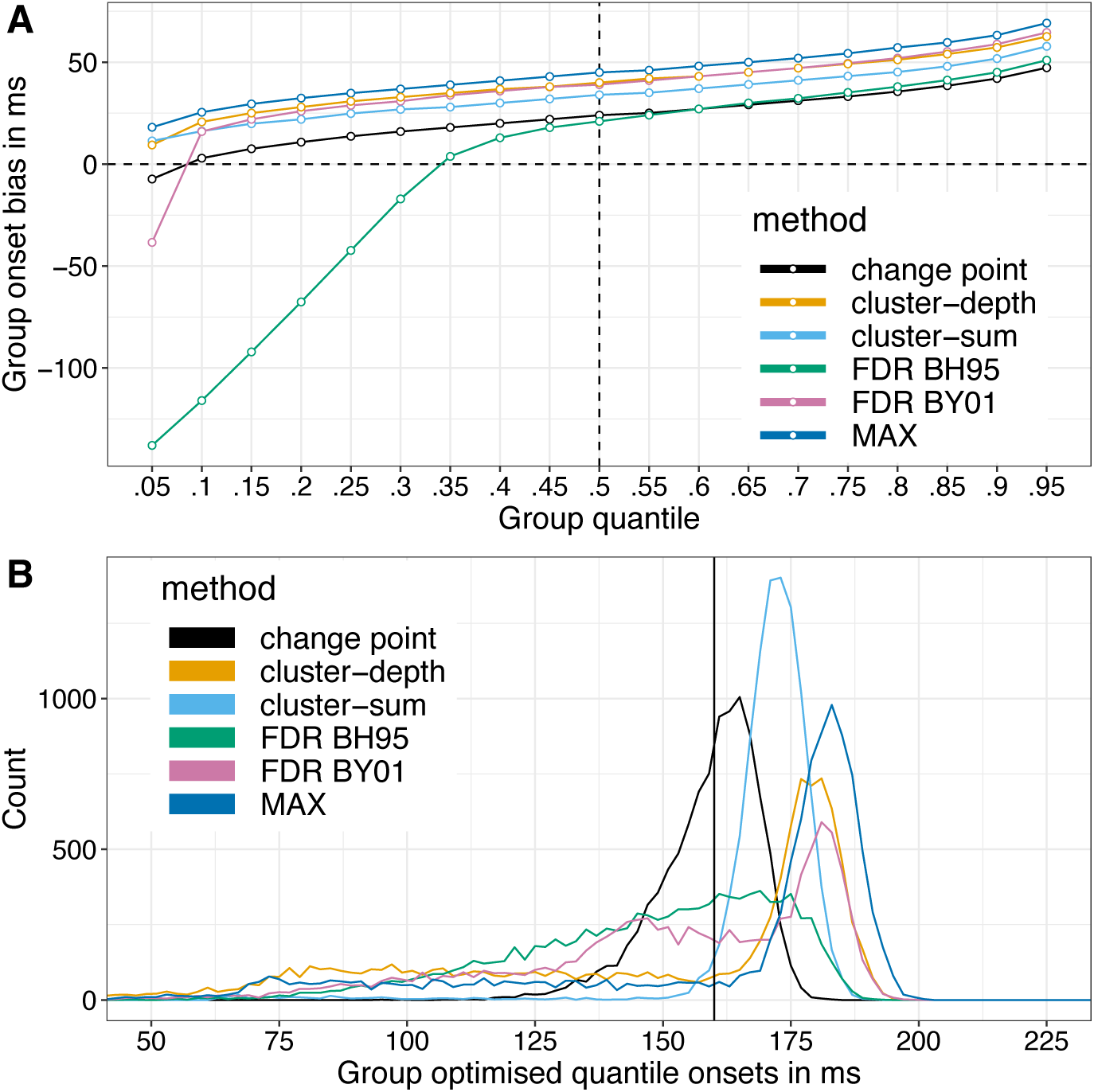
Group onset bias. (A) Group bias plotted as a function of group quantiles. The group onset quantiles were estimated using a median unbiased quantile estimator (Hyndman & Fan, 1996, definition 8). However, to handle tied values, there might be advantages in using the Harrell-Davis quantile estimator (Harrell & Davis, 1982; Wilcox & Rousselet, 2023). The vertical dashed line marks the median, leading to the results in Figure 7 and Table 2. (B) Optimised group onsets. Based on the results in panel A, matching sampling distributions are illustrated for the quantiles leading to the smallest positive bias: 0.1 quantile for BY01 and change point methods; 0.35 quantile for BH95; 0.05 quantile for the other methods. This figure can be reproduced by using the R notebook *onsetsim_eeg_group.Rmd*.

Given what we have learnt so far, what concretely could a group onset analysis look like? Imagine that we collected data from 50 participants, with 100 trials in each of two conditions. Data were simulated as in previous examples, with individual onsets drawn randomly from a 130-190 ms uniform distribution, and the maximum height of the signal drawn randomly from a 0.5-1 uniform distribution (arbitrary units).

Simulated individual mean ERP differences and the group average are illustrated in Figure 9A (first row). For each participant, point-wise t-tests were applied, and the resulting *t*^2^ values were used to estimate onsets using the change point algorithm (second row). Then, the 0.2 quantile of the distribution of individual onsets was used as a group estimate of the population onset (continuous vertical black line in all the plots in Figure 9). Finally, the measurement error associated with that group onset estimate was captured in a hierarchical bootstrap distribution (Rousselet et al. 2023): participants were sampled with replacement, then for each resampled participant, trials were sampled with replacement, independently from each condition, followed by point-wise t-tests and onset estimation using change point detection. Finally, the 0.2 quantile across all bootstrap participants was saved. The whole cascade of event was performed 2,000 times, leading to a distribution of 2,000 bootstrap estimates of the group onset (third row). This bootstrap distribution contains all the population values that are compatible with the data, given our model. The hierarchical bootstrap is a cheap way of integrating uncertainty across trials and participants, without an explicit hierarchical model. The bootstrap distribution can be summarised in different ways, for instance using a quantile interval, and here an arbitrary 97% quantile interval is illustrated (horizontal orange line in the third row of Figure 9).

**Figure 9.**
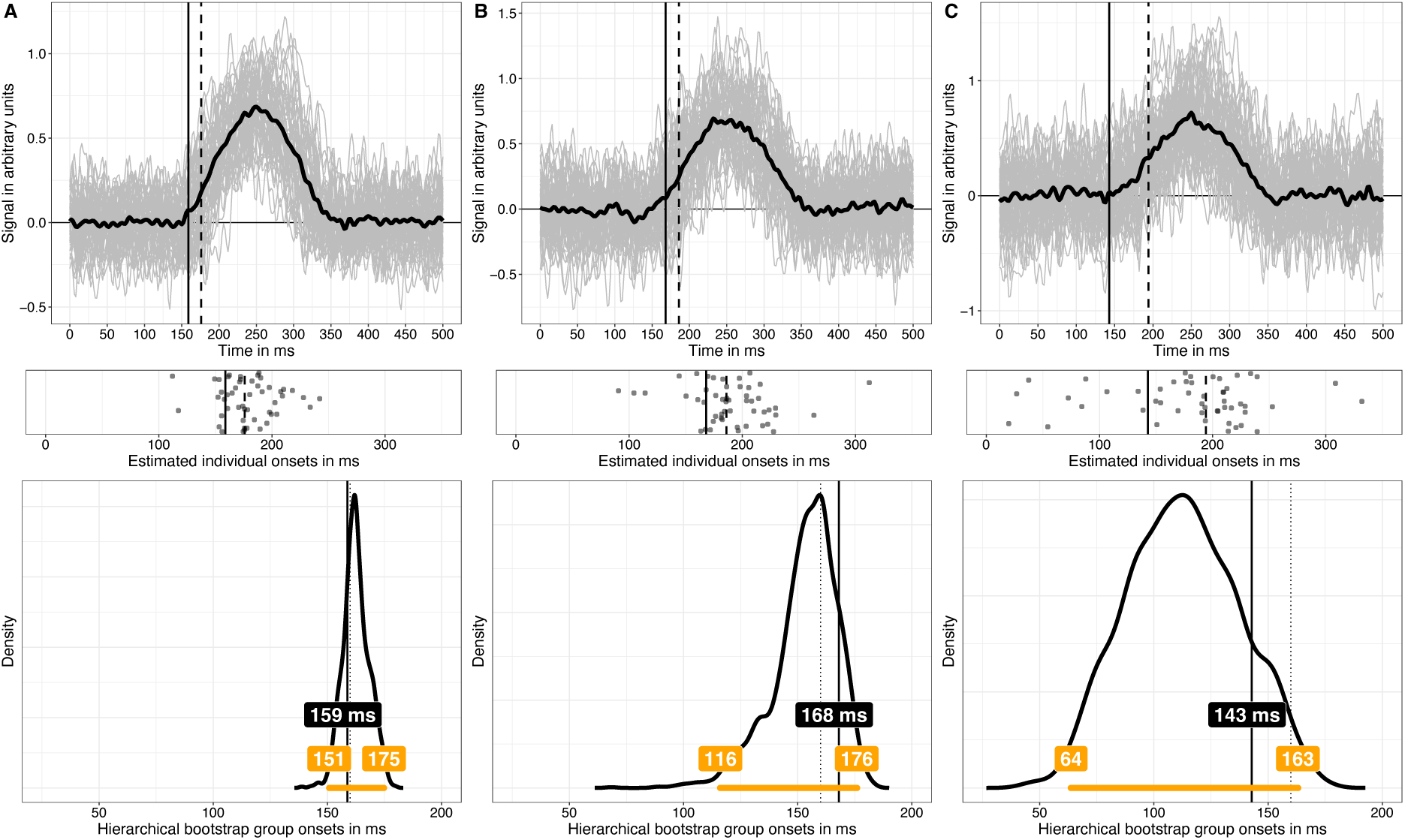
Simulated group data and hierarchical bootstrap analysis. In all columns, data from 50 participants were simulated, but the number of trials in each of two conditions varied: A=100 trials; B=50 trials; C=30 trials. Each panel in the top row shows in grey 50 individual time-courses of differences between the signal+noise condition and the noise only condition. The superimposed black time-course is the group average. The continuous vertical black line indicates the 0.2 quantile of the distribution of individual onsets; the dashed vertical line indicates the 0.5 quantile (median) of the same distribution. Individual onsets, estimated using the change point algorithm applied to *t*^2^ time-courses, are illustrated as scatterplots in the second row. The third row illustrates hierarchical bootstrap distributions of onsets. The continuous vertical line marks the 0.2 quantile onset. The dotted vertical line marks the population onset (160 ms). The horizontal orange line indicates a 97% hierarchical percentile bootstrap confidence interval, with the upper and lower bounds reported in the matching labels. The offset between the bootstrap distribution and the sample estimate of 0.2 quantile of onsets, particularly visible in column C, reflects that this quantity is biased, a topic covered extensively elsewhere (Rousselet & Wilcox, 2020). A bootstrap estimate of that bias is given by the difference between the estimate computed using the original sample and the mean or the median of the bootstrap estimates. This figure can be reproduced by using the R notebook *onsetsim_eeg_group_hb.Rmd*.

If instead of collecting 100 trials per condition, we only have 50 trials (Figure 9B), or 30 trials (Figure 9C), individual onset estimates are more spread out (second row) because individual time-courses are noisier, an increased uncertainty that is well captured by the bootstrap distributions. Similar results were obtained in a situation in which the number of trials is constant (n=50) and we vary the number of participants: 100, 50, and 30 (Figure 10).

**Figure 10.**
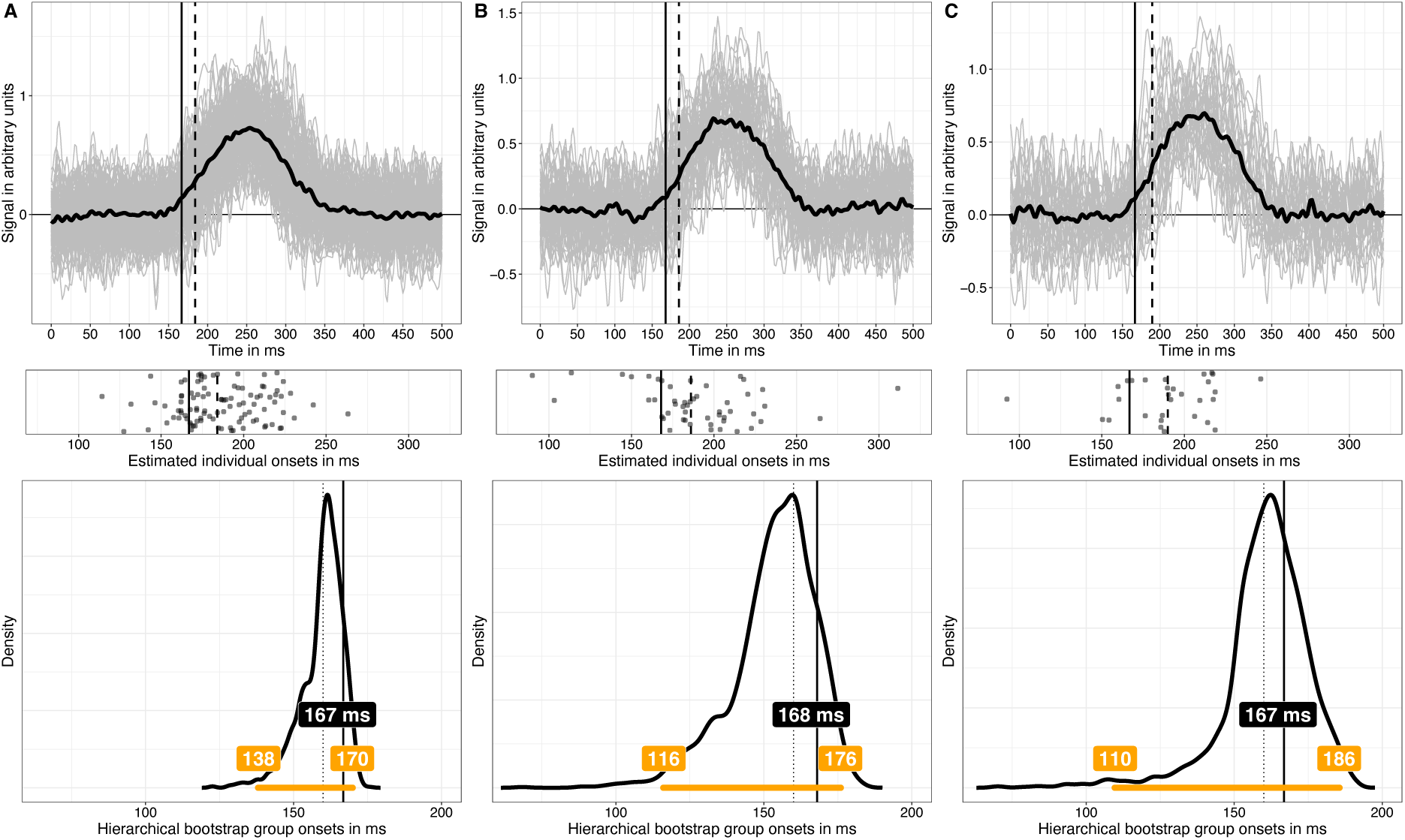
Simulated group data and hierarchical bootstrap analysis. Same as in Figure 9, but now the number of trials is constant (n=50), but the number of participants varied: A=100 participants; B=50 participants; C=30 participants. This figure can be reproduced by using the R notebook *onsetsim_eeg_group_hb.Rmd*.

To finish, it is worth noting another simple way to tackle onset estimation bias: group comparison. If onsets are estimated in two independent groups of participants, and then compared, the bias in each group will be washed away in the comparison, if the data in the two groups lead to similar biases. The assumption of equal bias between groups might not hold for instance if they differ in ERP shape or signal to noise ratio. The interpretation of such comparisons also requires the strong assumption of linear mapping between what is measured (say ERP onset) and the target of investigation (say processing speed), a consideration not often seen in applied research (Kellen et al., 2021; Wagenmakers et al., 2012). If we are willing to make these assumptions, a simple simulation can demonstrate the lack of bias in group comparisons. Imagine that we have two groups, each containing 20 participants. Participant level data are generated as in previous examples, with 50 trials in each of two conditions. Onsets are estimated by applying change point detection to *t*^2^ time-courses, and the means of the two groups are compared. In the first group of participants, onsets vary uniformly between 140-180 ms. In the second group of participants, we consider three situations: [1] onsets are also distributed uniformly 140-180 ms (no difference); [2] onsets are distributed uniformly 130-190 ms (variance difference); [3] onsets are distributed uniformly 150-190 ms (10 ms shift). Figure 11 illustrates the sampling distributions of 10,000 simulated group differences for the 3 types of group comparisons. The distributions are symmetric, as expected, and show no sign of bias. Finally, if the two group distributions of onsets differ in shape, a likely prospect, a more powerful approach is to compare the groups at multiple quantiles instead of a single measure of central tendency (Bieniek et al., 2016; Rousselet et al., 2017; Wilcox & Rousselet, 2024).

**Figure 11.**
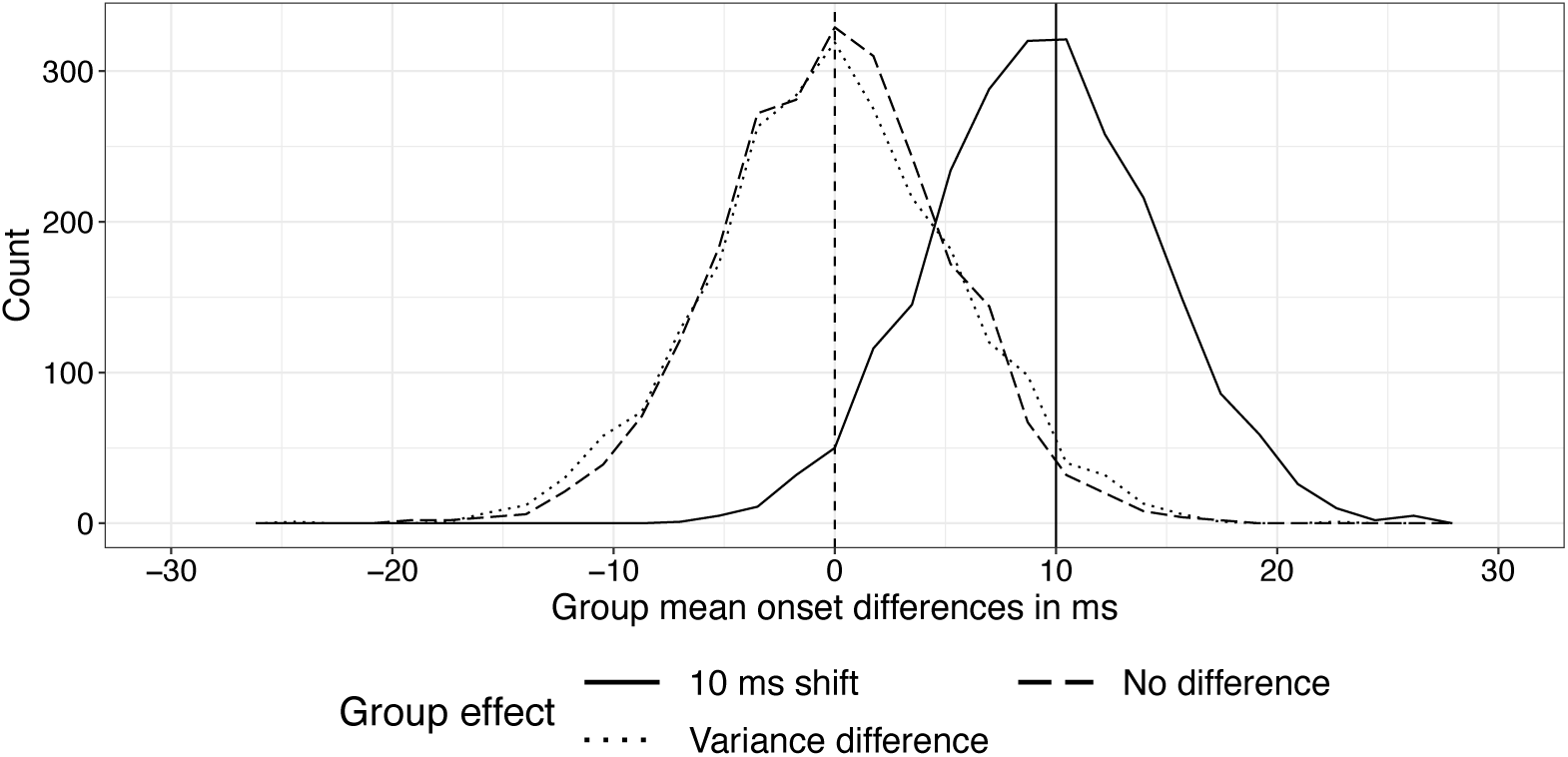
Sampling distributions of group onset differences. Each distribution consists in 10,000 onset differences between the means of two independent groups. For each comparison, a reference group of 20 participants was compared to another group that was sampled either from the same population (no difference), a population with more onset variability (variance difference) or a population with later onsets (10 ms shift). This figure can be reproduced by using the R notebook *onsetsim_eeg_group_comp.Rmd*.

## 4. Discussion

Onsets are inherently noisy because they correspond to the testing points with the lowest signal to noise ratio. As a result, we can expect most onset estimation methods to produce positively biased and variable results. Consistent with that intuition, Sassenhagen & Draschkow (2019) demonstrated that the cluster-sum method to correct for multiple comparisons in a mass-univariate approach can lead to high variance and positively biased onset estimation. The simulation results presented here confirm their results. However, comparing cluster-sum to other methods helps put the results in perspective: the cluster-sum approach performs relatively well compared to other standard methods, in terms of bias and measures of variability. Certainly, users interested in measuring onsets will achieve dramatically less variable results by using cluster-sum than standard FDR methods. Whether more recent FDR methods, including variants developed specifically for ERP data, can perform better remains to be determined (Korthauer et al., 2019; Sheu et al., 2016). My preliminary investigation suggests that local FDR and *q* values (Strimmer, 2008) do not improve performance over BH95 and BY01. Users of the cluster-sum approach will also achieve less biased results relative to the more conservative MAX and cluster-depth methods.

Despite the relatively good performance of cluster-sum in the simulations, it remains problematic if the goal is to achieve point-wise error control. Indeed, as reviewed in the introduction, techniques that provide weak FWER control cannot be used to make claims about specific time-points, including the location of onsets. But the issue of error control for onsets is not solved by standard methods that provide strong FWER control either: whatever the type of control afforded, and whether point-wise *p* values are available, none of the common mass univariate techniques in EEG and MEG were designed to estimate onsets. Using these methods for onset estimation necessarily involves several statistical errors, most notably an interaction fallacy, because the implied interaction between a segment with a series of non-significant tests and a segment with a series of significant ones is never explicitly tested (Nieuwenhuis et al., 2011).

In that respect, a promising approach is to give up entirely on the mass-univariate approach to estimate onsets, and instead employ change point detection algorithms, which have been developed explicitly for that purpose. Whether they work well with EEG and MEG data is an empirical question; the simulation results presented here are certainly encouraging, especially given that I used a relatively old algorithm. More recent algorithms might perform even better (Aminikhanghahi & Cook, 2017; Lindeløv, 2023; Truong et al., 2020), including Bayesian approaches that allow hierarchical modelling (Lindeløv, 2020). However, there is a catch: the simulations relied on the presence of clear effects. Applying the binary segmentation to *t*^2^ time-courses for ERP conditions that do not differ would lead to spurious onsets. So, instead of abandoning mass-univariate approaches, we could see a division of labour: apply a mass-univariate method in exploratory work to establish the presence of an effect in individual participants—because of its lower computational cost, the MAX approach would be particularly useful here. Then apply a change point detection method when an effect is detected. Alternatively, change point detection could be better calibrated (Truong et al., 2020), or it could be applied to less noisy metrics, such as mutual information (Ince et al., 2017). Whatever the method, the resulting individual onsets can then be used to make group inferences, for instance to assess how the timing of an effect differ between groups or conditions, similarly to the much more common analyses of peak amplitudes and latencies.

Such hierarchical approach, involving single-participant estimation followed by group inferences is not the norm but is nevertheless well established (for instance Pernet, Sajda, et al., 2011; Rousselet et al., 2011; Rousselet & Pernet, 2011). In contrast, typical mass-univariate ERP group analyses forfeit individual differences by only reporting statistical significance at the group level, leading to a single onset being reported, without any measure of uncertainty. That practice would be considered unacceptable for behavioural data, such as reaction times or percent correct data. Why it is deemed acceptable for ERP effects is unclear, especially when there is so much to gain from inferences at the participant level (Ince et al., 2017, 2021). For instance, combined with quantile inferences, the hierarchical approach can be used to test more specific hypotheses about the timing of neuronal events (Bieniek et al., 2016; Rousselet et al., 2017). As demonstrated in Figure 8, using lower group quantiles can also reduce group estimation bias, though this will require simulations for specific applications, for ERP effects that differ in strength, duration as well as within- and between-participant variability. In particular, the examples presented in Figures 9-10 raise the issue of how to spend our resources: investing in more trials or in more participants (Rouder & Haaf, 2018)? Depending on the nature of the effect, our goals and cost functions, there will be more to gain in adding one or the other.

In addition to increasing sample sizes, other useful steps could be considered to reduce bias and increase estimation precision, such as robust estimation, instead of relying on mean estimation by default (Ince et al., 2017; Wilcox & Rousselet, 2023). At the pre-processing stage, filtering characteristics should be considered carefully, with non-causal high-pass filters introducing potentially severe signal distortions (Rousselet, 2012; van Driel et al., 2021; Widmann et al., 2015; Widmann & Schröger, 2012). Instead, causal filters can be very effective at attenuating low temporal frequencies while avoiding backpropagation of effects, both of which can distort onsets. Using data spatially deblurred could also improve onset estimation, for instance by working with independent components or current source densities (Delorme & Makeig, 2004; Tenke & Kayser, 2012). On the stimulus front, screen luminance can strongly influence onsets and should be considered when comparing results between studies (Bieniek et al., 2013). It is also very informative to assess the test-retest reliability of onsets, either using the same data (via cross-validation), or data from a different EEG session. For instance, test-retest assessment of face ERP onsets suggest that the individual variability in onsets is relatively stable across sessions, indicating that individual differences reflect more than measurement noise (Bieniek et al., 2016). Further, computational consistency can be assessed by reporting results from more than one onset estimation method.

Finally, it is worth considering how the analysis pipeline proposed here as a proof of concept could be applied to multiple electrodes. A simple solution could be to apply a change point detection method independently at each electrode, and then summarise the onset distribution using a calibrated quantile, based on simulations. A second option would be to perform change point detection on a virtual electrode, which could be obtained by using the maximum statistics across electrodes (Jaworska et al., 2020), or by working in source space, for instance on independent components (Delorme et al., 2012). There are also change point algorithms that work on multivariate time-series (Grundy et al., 2020; Moradi et al., 2023). There is exciting simulation work to be done on all these topics.

## ABBREVIATIONS

MEG: magnetoencephalography
EEG: electroencephalography
ERP: event-related potentials
fMRI: functional magnetic resonance imaging
FWER: family-wise error rate
FDR: false discovery rate
MAX: maximum statistics
BY01: Benjamini & Yekutieli (2001)
BH95: Benjamini & Hochberg (1995)
CS: cluster-sum algorithm
CD: cluster-depth algorithm
CP: change point algorithm
MAE: mean absolute error
ms: millisecond

## Data and code availability

The R and Matlab code necessary to reproduce the simulations is available on GitHub: https://github.com/GRousselet/onsetsim. The repository also contains simulation results and code to reproduce the figures in this article, without running the simulations. The main R packages I used include ggplot2 (Wickham, 2016), cowplot (Wilke, 2017), Rfast (Papadakis et al., 2019), changepoint (Killick & Eckley, 2014), permuco (Frossard & Renaud, 2021), and the essential beepr (Bååth, 2018).

## Author’s contributions

**Guillaume Rousselet:** conceptualisation, formal analysis, methodology, software, visualisation, writing.

## Acknowledgements

I thank Jona Sassenhagen, Benedikt Ehinger, and two anonymous reviewers for their very useful comments on an earlier version of this article. In addition to reviewing the article, Dr Ehinger also reproduced the main simulations, using his own implementation in Julia (Ehinger, 2024).

